# Towards decoding inner speech from EEG and MEG

**DOI:** 10.1101/2025.10.13.682161

**Authors:** Richard Csaky, Mats W.J. van Es, Oiwi Parker Jones, Mark Woolrich

## Abstract

Despite the prevalence of inner speech in everyday life, research on this has been limited, particularly when it comes to non-invasive methods. This preprint aims to fill this gap by using EEG and MEG to collect data from three different inner speech paradigms, and by conducting an initial decoding analysis. Specifically, we tested silent reading, repetitive inner speech, and generative inner speech tasks.

We collect a high number of inner speech trials from a few participants. Besides comparing across recording modalities we also compare across inner speech types. Our aim is to analyse the decodability of inner speech within each task and between tasks by the use of transfer learning. We find that in both EEG and MEG, silent reading can be decoded relatively well with 30-40% accuracy across 5 words. However, the decoding performance of both types of inner speech is mostly at chance level. This prohibited further transfer learning investigations between tasks. While the inner speech results are primarily negative, we believe our exploration of data size and various decoding methods is valuable. The dataset itself is useful for the research community as it contains a much larger number of trials within one participant than any other inner speech dataset. Having multiple sessions also allows for testing across-session performance.

Finally, we systematically compare silent reading decoding performance within 3 participants across four non-invasive modalities. These are EEG, 2 types of MEG machines, Elekta and CTF, and optically-pumped magnetometers (OPMs). We also compare the spatiotemporal dynamics of silent reading between these modalities. This is especially aimed at validating OPMs as a new kind of non-invasive brain recording technology. We find comparable performance to EEG, but OPM performance did not reach traditional MEG.

## 1 Introduction

Inner speech, also known as verbal thinking or covert self-talk, refers to the internal monologue that occurs within one’s mind. This phenomenon has been extensively studied in psychology and cognitive science (Alderson-Day and Fernyhough, 2015). More recently, neuroimaging techniques have allowed researchers to examine the neural correlates of inner speech directly. As discussed by Geva (2018), early 20th century studies used measurements of tiny muscle movements during imagined speech production to infer inner speech. However, the advent of modern neuroimaging techniques like MEG and fMRI has enabled more direct investigation of the brain regions involved in inner speech compared to overt speech.

Traditional brain-computer interface (BCI) systems are relatively slow and do not leverage inner speech, which has the potential to enable communication at natural speech rates. Some progress has been made in decoding visual stimuli during reading limited sets of words or sentences (Mugler et al., 2014; Hultén et al., 2021; Moses et al., 2019), as well as decoding speech perception and overt speech production where muscle movements are present (Dash et al., 2020b; Défossez et al., 2022). However, detecting brain signals associated specifically with inner speech remains challenging given the lack of external stimuli or produced behaviour to provide timing information.

While we focus here on the hard problem of non-invasive inner speech decoding, more tractable approaches exist. These include applying invasive methods to inner speech, as well as applying non-invasive methods to related tasks that share features with inner speech. Related tasks either provide external timing information through stimuli like silent reading/listening, or leverage produced behaviour like overt loud or silent speech. However, it is unclear how insights from these tasks might transfer to decoding inner speech itself. Disentangling task-related activity (e.g. visual, auditory, or motor) from pure inner speech is difficult. Similarly, models tuned to detect muscle activation during silent speech may not transfer well to pure inner speech decoding (Dash et al., 2020b).

There are also experimental challenges inherent to studying inner speech. While people often experience spontaneous inner speech during mind wandering, there is no way to align such endogenous thought with recorded brain activity. This leaves two options: a purely unsupervised learning approach, or using more constrained, artificial experiments. The latter often employs visual or auditory cues to elicit inner speech in a time-locked manner. Some papers refer to cued silent reading tasks as a form of inner speech. However, we argue this confounds inner speech with concurrent visual processing of the words.

Pure inner speech paradigms can be achieved by using identical cues across different conditions. Brain responses to the cues themselves will be consistent, while activity varying between inner speech conditions can be disentangled. A limitation is that using identical cues provides no overt record of the specific condition of each trial. Participants can either be instructed on the inner speech to generate for each cue beforehand (*repetitive* inner speech), or report the contents after each trial (*generative* inner speech) (Parker Jones and Voets, 2021). Both approaches enable studying inner speech independently from external stimuli or behaviour.

The aims here are multifaceted, but guided by advancing non-invasive BCI communication. We approach this challenge through comparing:

1. Non-invasive recording modalities: EEG, MEG, OPM.
2. Types of inner speech: silent reading, repetitive, generative.
3. Data quantities: number of trials and sessions.
4. Decoding methods.

Our focus is on enabling BCI applications rather than basic cognitive neuroscience of inner speech per se. We review relevant research on decoding inner speech using invasive and non-invasive neural recording methods in Section 4. Next, we describe our experimental paradigms and analysis methods for investigating non-invasive inner speech decoding.

## 2 Methods

Our experimental paradigm follows previous efforts to delineate repetitive and generative inner speech. In a similar line of work Parker Jones and Voets (2021) investigate whether neural decoders trained on elicited inner speech data can be successfully transferred to decode self-generated inner speech (see Figure 1 for task paradigms). Using fMRI data from one subject, the authors trained deep neural networks on a large dataset of elicited inner speech collected during covert reading and repeating tasks. They then tested these models on new self-generated inner speech data collected while the subject freely imagined syllables. The transferred decoders predicted unseen phonemes with high accuracy. The successful zero-shot task transfer demonstrates the viability of leveraging elicited speech to train models that can decode self-generated inner speech. This has practical significance for developing inner speech brain-computer interfaces, since elicited tasks allow collection of labelled training data even from locked-in patients.

**Figure 1:**
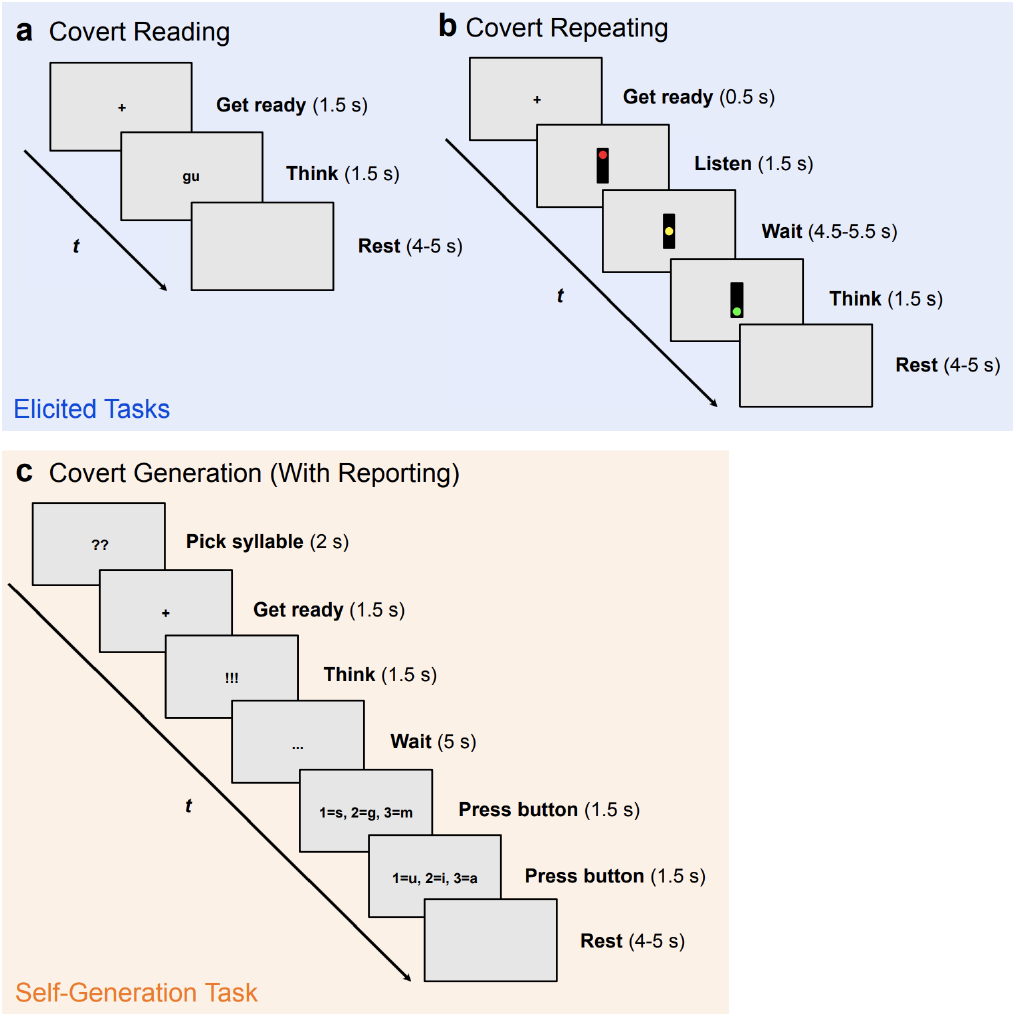
Visualisation of silent reading (a), repetitive (b) and generative inner speech (c) paradigms used in Parker Jones and Voets (2021). Figure from Parker Jones and Voets (2021).

While previous studies have investigated various language units such as characters, phonemes, words, and even whole phrases, we focused our experiments on the word level. A limited set of words already has direct benefits for potential patients. Working with words also allows scaling up to cover the entire language lexicon in future research. Since we are interested in potential clinical applications, we chose a set of words that could be most useful for patients: help, hungry, tired, pain, thirsty. Decoding at the individual rather than group level is critical for practical applications. Thus we conducted our experiments with a small number of participants. Selecting only 5 words also allows us to test how collecting a large number of trials per word can improve decoding performance.

We collected data across two versions of the experimental paradigm, adapting it as we gained results to better align with our objectives. In version 1, we collected 1-second silent reading trials followed by 4 consecutive 1-second repetitive and generative inner speech trials. We collected a small MEG dataset, concentrating efforts on obtaining a high number of EEG sessions from the same participant to assess between-session transferability. In version 2 we omitted inner speech entirely and collected only silent reading data across 4 modalities - two MEG systems (Elekta and CTF^1^), EEG, and optically pumped magnetometers (OPMs). In version 1, we used an Elekta Neuromag Triux 306-channel system for MEG scans. We used a Neuroscan 64-channel cap for standalone EEG (standard 10-20 layout), with MEG-compatible Easycap EEG used for combined MEG and EEG.

For combined M/EEG, EEG ground and reference were on the left cheek and nose, respectively. For standalone EEG, the Cz and POz locations served as reference and ground, and we placed extra electrodes on the two mastoids. These could also serve as reference in offline analysis. Voltage and thus signal is always measured relative to a reference electrode in EEG. This means that the signal is the difference in voltage between the reference and other electrodes. Thus, the choice of reference greatly influences the characteristics of the recorded signal. It is best practice to place the reference electrode on the head but away from locations which might contain signals of interest. In our case both reference placements satisfy these criteria. While the signal shape and evoked responses will look different with different reference choices, this does not matter for decoding applications, as long as the signal of interest is not accidentally removed.

For most scans, we simultaneously collected electrooculogram (EOG) and electrocardiogram (ECG) data for easier artifact removal. ECG electrodes were placed on the wrists, with horizontal EOG on the outer side of the eyes and vertical EOG above and below the left eye. Electromyography (EMG) electrodes on the jaw monitored subtle mouth movements. Structural MRI scans were obtained for all participants in versions 1. During Elekta scans, we video recorded the mouth of participants to ensure no task-related motion. Eye tracking was performed for all MEG and EEG scans. We collected 5 minutes of resting state data before and after each scan. Stimuli were delivered via PsychoPy. The experiment was reviewed and approved by the Medical Sciences interdivisional research ethics committee at the University of Oxford (reference number R75957/RE001).

### 2.1 Experimental paradigm

**Version 1** In the first version of the experiment, participants silently read words displayed individually on a screen, followed by 4 consecutive visual fixation-cross cues to covertly repeat the word (Figure 2). This phase lasted approximately 5 minutes. In the next phase, participants continued reading and repeating words, but were now prompted after 0-2 read/repeat trials to imagine speaking a different word from the 5-word set (the generative inner speech task). 0-2 means that we randomly sampled either 0, 1, or 2 consecutive read and repeat trials. Similarly to the repetitive task we prompted the inner speech with 4 consecutive visual fixation-cross cues. Participants then indicated their imagined word with a button press. The random 0-2 read/repeat trials before each generative prompt ensured that participants did not pre-select words, better resembling unconstrained inner speech. We noticed that pure generative blocks let participants pre-plan words upon indicating the previous selection. Introducing random read/repeat trials limited this behaviour.

**Figure 2:**
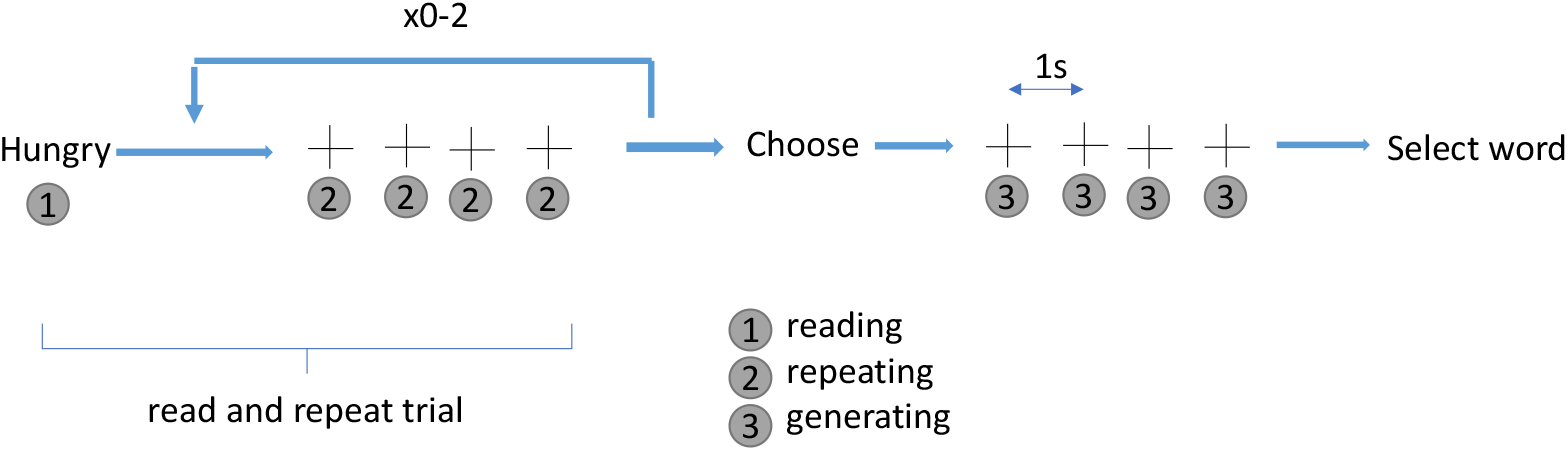
Paradigm for version 1 of our experiments. The participant silently *reads* ‘Hungry’, then *repeats* it four times at 1-second intervals cued by crosses. This can repeat 0-2 times before *generating* a new word from the 5-word set at four 1-second cross cues, avoiding the previous read/repeat word(s).

Each cross was displayed for 0.3 seconds followed by a 0.7 second-long blank screen. Word stimuli were displayed for 0.8-1.0 seconds, followed by a 0.8-1.0 second-long blank screen. Word order was randomised. The total second phase duration was approximately 50 minutes in 4 blocks with breaks between blocks. We collected simultaneous MEG and EEG data at the Oxford Centre for Human Brain Activity (OHBA). While we collected some combined MEG, we focused on obtaining multiple EEG-only sessions from one participant to assess between-session transferability.

**Version 2** As the inner speech tasks yielded poor results, the second version focused solely on silent reading trials to collect more data within the 1-hour sessions. We expanded our aims and collected combined M/EEG at OHBA along with CTF-MEG and OPM data (same participants) in collaboration with the OPM Lab led by Matthew Brookes at the University of Nottingham. The simple paradigm displayed the 5 words randomly for 0.8-1.0 seconds with 0.8-1.0 second breaks. After every 10 trials, participants indicated the last word read to monitor attention.

### 2.2 Analysis

#### Data acquisition and preprocessing

Elekta and EEG data were acquired at a sampling rate of 1000 Hz with a built-in bandpass filter between 0.03 and 330 Hz, while CTF and OPM scans were acquired at a sampling rate of 1200 Hz with a built-in lowpass filter of 600 Hz. The Elekta system contained 102 magnetometers and 204 planar gradiometers (102 sensors x 2 orientations), totalling 306 channels. CTF contained 265 axial gradiometers. Sensor configurations for three modalities are illustrated in Figure 3. OPM data were recorded using a variable number of triaxial magnetometers (typically 150-180) in 60 fixed scalp locations, measuring magnetic fields along orthogonal axes. Channel configurations for individual participants are reported in Section 3.1.

**Figure 3:**
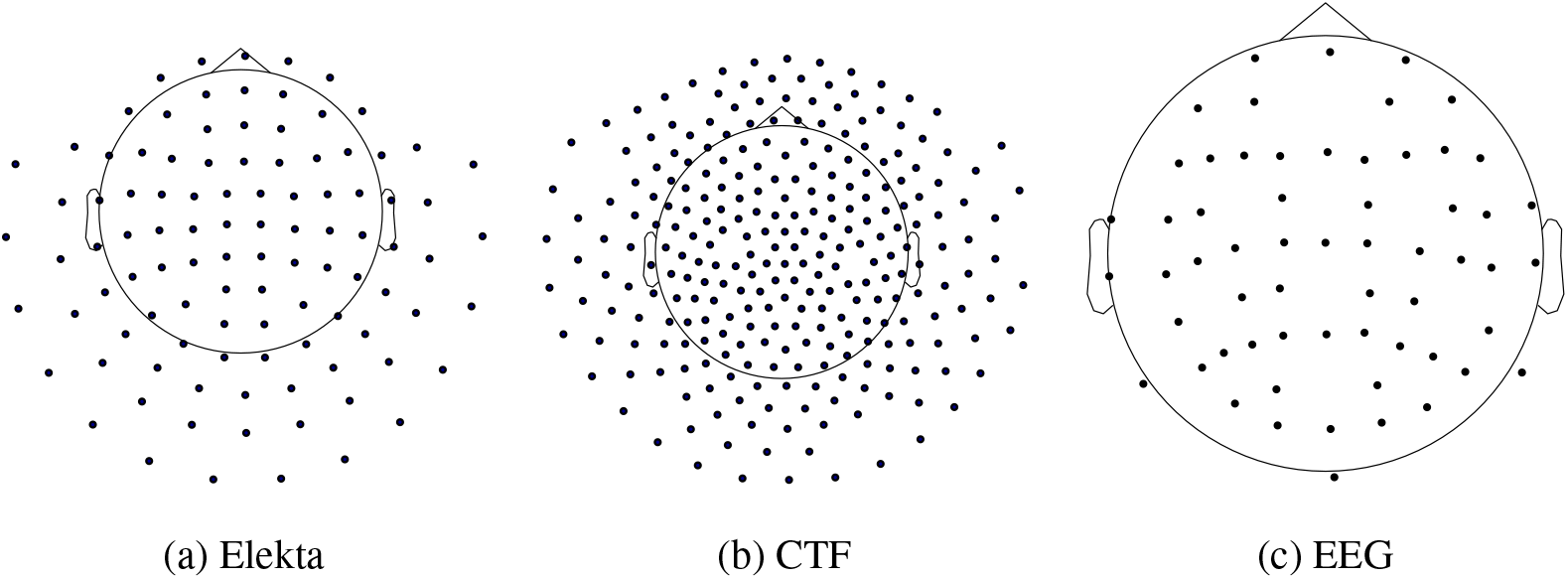
Sensor locations across scanning systems. CTF contained 1 gradiometer per location, while Elekta had 2 gradiometers and 1 magnetometer. Please note that OPM sensor layouts are reported in Section 3.1.

Elekta data were preprocessed using Maxwell filtering for movement compensation and signal space separation using the MaxFilter algorithm (Taulu and Simola, 2006). Noisy OPM channels identified during recording were removed prior to analysis. For all systems, data were bandpass filtered (1-25 Hz typically, with higher lowpass for some experiments), notch filtered at 50 and 100 Hz, and subjected to automated bad channel detection (except OPM) using the oslpy package^2^ (Quinn et al., 2022). Bad segments were identified via a multi-pass procedure across progressively wider temporal windows (200 to 800 ms) with a significance threshold of 0.1. Independent component analysis was applied for dimensionality reduction (64 components for MEG/OPM, 32 for EEG). Components reflecting ocular or cardiac artifacts were removed before downsampling to 100 Hz and epoching.

An additional mean field correction was applied to OPM data after preprocessing:

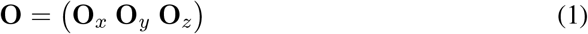

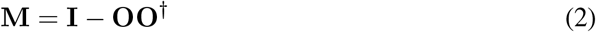

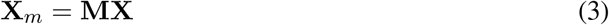

Where **O**_*x*_, **O**_*y*_, **O**_*z*_ are sensor orientation vectors, **O** stacks them vertically, **I** is the identity matrix, and **M** is the final mixing matrix applied to the data **X**. The aim of this transformation is to remove any spatially homogeneous field not coming from the brain and is described in detail in Tierney et al. (2021). Elekta data already includes such corrections in the MaxFilter algorithm, and CTFs do not need it as they measure the field gradient with gradiometers only.

Basic analyses involved comparison of evoked responses across channels, conditions, sessions, and modalities. Single-trial covariance matrices were also computed and visualised using t-SNE (Maaten and Hinton, 2008).

#### Decoding

We employed several decoding methods for both the silent reading and inner speech tasks. For inner speech we tried several methods, but mainly used the covariance matrix over each trial as features for an LDA model. Channels were standardised prior to decoding. We tried concatenating across the 4 consecutive trials to form 4-second epochs, as well as averaging over trials, before decoding. Specific methods and hyperparameters are detailed in the Results.

## 3 Results

### 3.1 Data statistics

A total of 4 male participants (P2, P4, P5, P6) between the ages of 20 and 40 participated across the two versions of our study. The participant pool included both native and non-native English speakers, though all had C2 level proficiency in English. Participant 4 (P4) is the author of this thesis and a non-native English speaker.

The two versions of the experiment were conducted with different goals. Version 1 involved evaluating the feasibility of decoding inner speech in 3 participants. Since EEG and MEG provided comparable decoding accuracy, 10 EEG sessions were collected from P4 in version 1 to examine improvements from increased data size and test decoder adaptability across sessions. In version 2, the inner speech task was removed and silent reading trials were collected using combined M/EEG, CTF, and OPMs across 3 participants. P4 and P5 also participated in version 1. The high number of silent reading trials (1250 per session) enabled thorough investigation of this paradigm. The OPM sensor layouts are shown in Figure 4.

**Figure 4:**
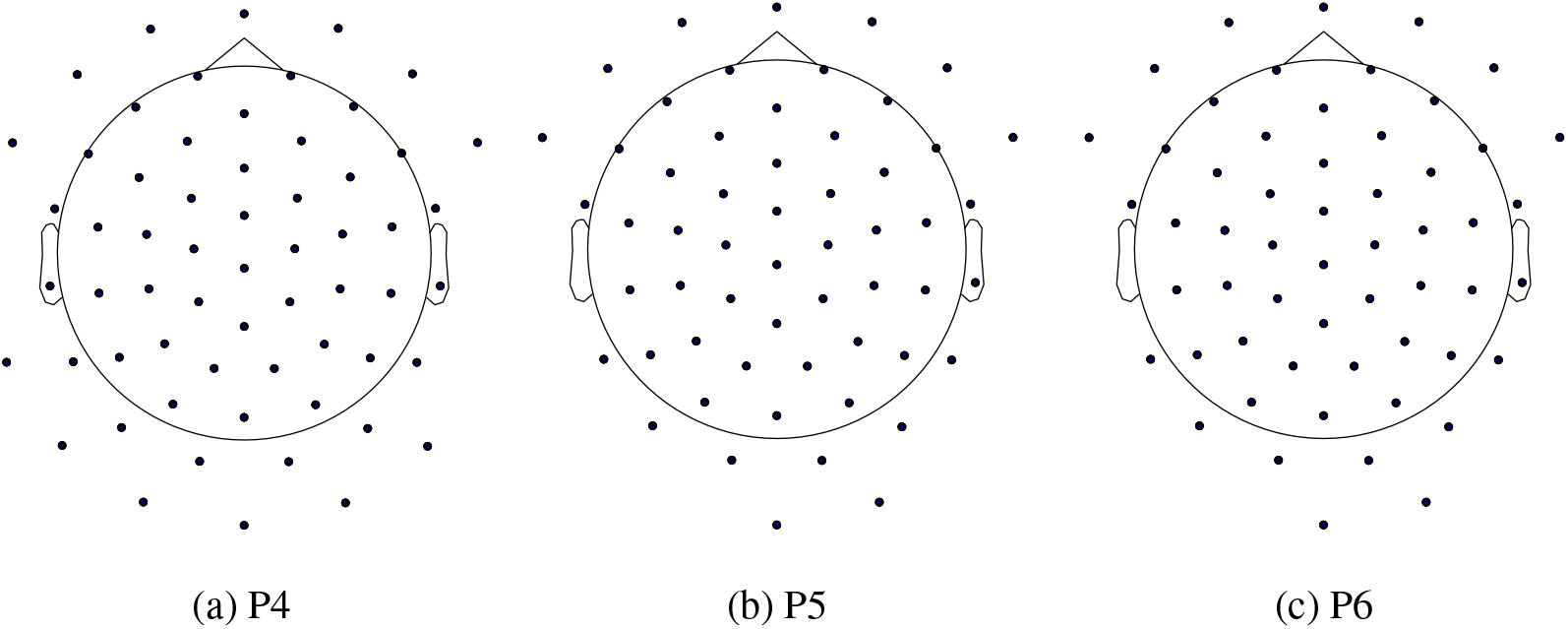
OPM sensor configurations across the three participants in version 2 of the experiment. Each location contained an OPM sensor measuring the magnetic field in three orthogonal directions. Sensor layouts and number of sensors are different due to technical difficulties with operating all sensors without overheating, excessive noise, or other issues.

Tables 1 and 2 summarise the number of sessions and trials for the participants in each version.

**Table 1:**
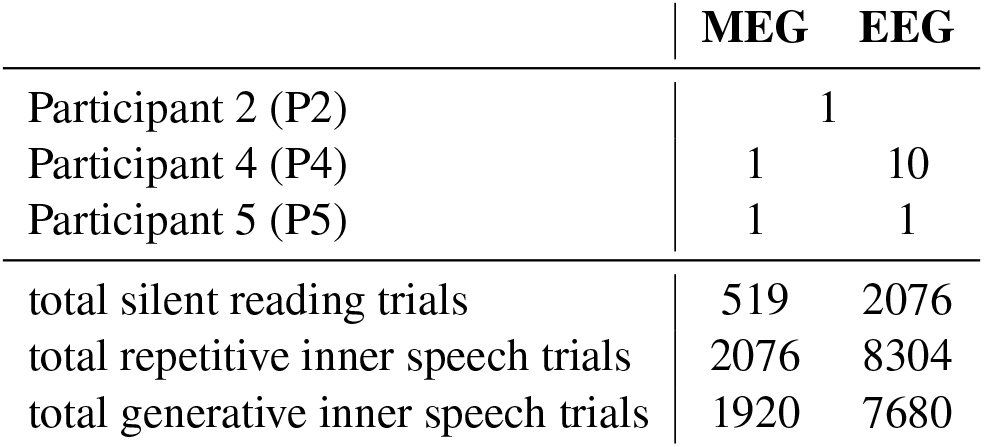
Number of sessions for each participant in version 1 of the study (top 3 rows). Total number of trials is given across all sessions and participants. Number of trials may be slightly lower or higher than shown due to randomness. Note that for P2 we conducted a combined M/EEG session while for the other participants MEG and EEG scans were separate.

**Table 2:** Number of sessions for each participant in version 2 of the study (top 3 rows). Total number of trials is given across all sessions and participants.

|  | combined M/EEG | CTF | OPM |
| --- | --- | --- | --- |
| Participant 4 (P4) | 1 | 1 | 1 |
| Participant 5 (P5) | 1 | 1 | 1 |
| Participant 6 (P6) | 1 | 1 | 1 |
| total silent reading trials | 3750 | 3750 | 3750 |

While target numbers of trials are reported, minor variations occurred due to randomisation. The extensive datasets collected enabled thorough investigation of silent reading and inner speech paradigms. The multiple sessions also allow examination of between-session and between-modality variability. Overall, the dataset provides a unique resource to advance decoding of covert speech from non-invasive electrophysiological signals.

### 3.2 Data analysis

In this section we present our non-decoding analyses of the collected data. The aim of this analysis is to validate data quality and uncover any insights into differences between tasks. These visualisations were primarily conducted on the 10 EEG sessions of P4, as this participant had the most inner speech trials (version 1 of the study) and sessions, allowing investigations into between-session variability. For the visualisations in this section, no independent component analysis (ICA) artifact removal was performed on the data. We plot the electrode positions and their names over the scalp in Figure 5.

**Figure 5:**
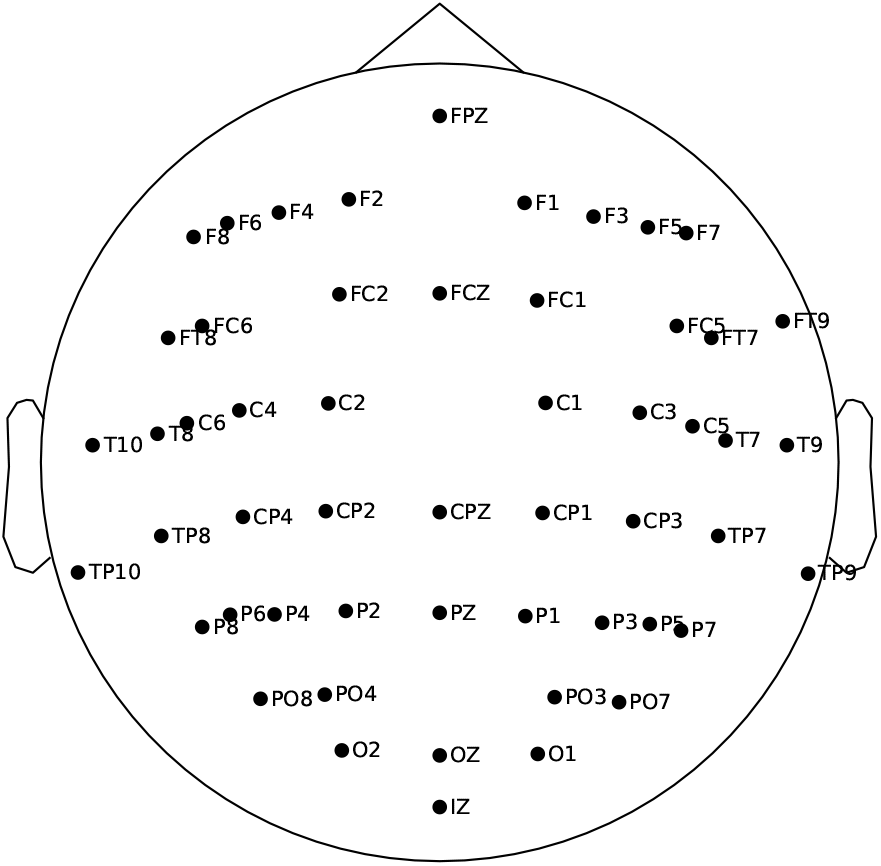
EEG electrode locations for P4 in version 1 of the experiment.

To visualise between-session variability, we compute the covariance across each 1-second inner speech trial, and average these across individual sessions. Figure 6 exhibits the averaged covariance of each session. Channels demonstrate high covariance within brain regions, such as the frontal and visual areas. Across sessions, average covariances appear similar. To quantify similarity between sessions, we computed the Riemannian distances of the covariances between pairs of sessions for all possible pairs. This produces a session-by-session distance matrix (Figure 7). This can provide insight into between-session differences. For instance, the first session seems quite distant from the other sessions. This can mean that a decoder trained on other sessions may not perform very well on this session.

**Figure 6:**
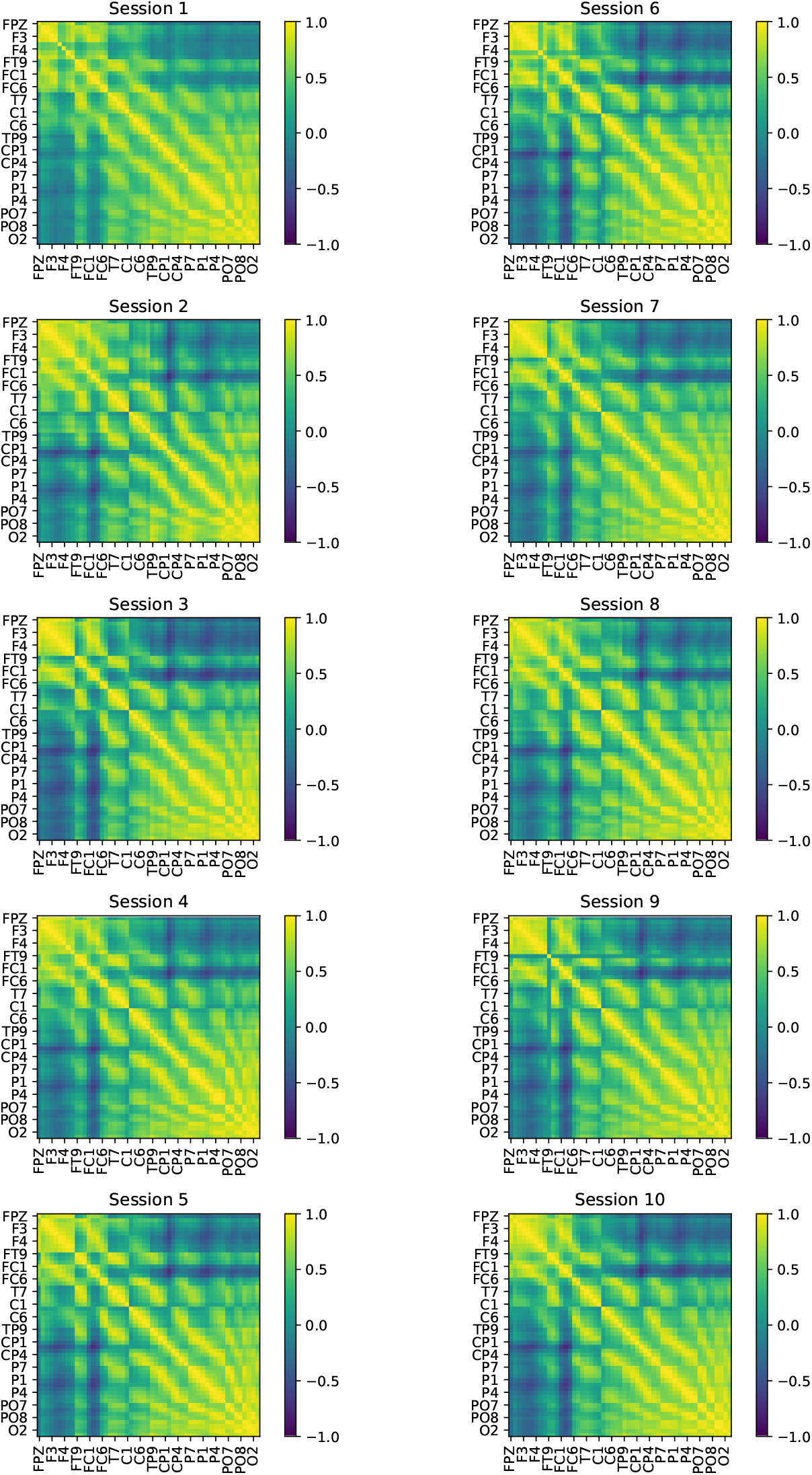
Averaged trial-covariances across the 10 EEG sessions of P4 in version 1 of the experiment. Each matrix represents a different session.

**Figure 7:**
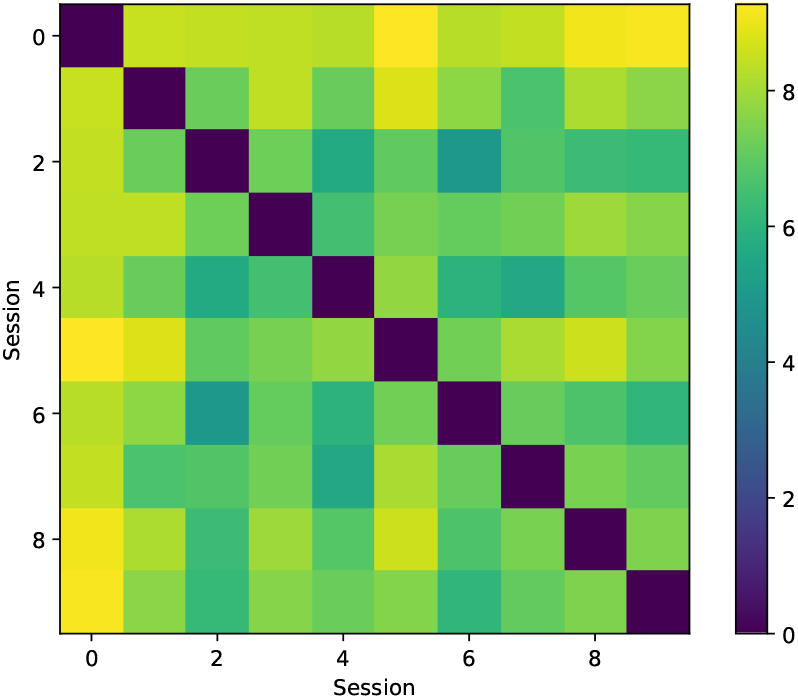
Riemann distance matrix between the average session-covariances across the 10 EEG sessions of P4 in version 1.

Finally, we investigated whether the covariance representations demonstrate interesting structure when visualised in 2D. To this end, we simply applied t-SNE to the individual trial covariances to project them into 2 dimensions and visualised the result. Figure 8 portrays this projection with two types of labelling. When trials are labelled by their corresponding condition (word), no apparent clustering emerges. This further bolsters that differentiating between words in inner speech is challenging. Structure can be discerned in the projection when labelled by session. This is anticipated since trials within one session are more similar to each other than to other sessions. These findings imply that decoding inner speech may be an equally challenging endeavour. An investigation of evoked responses is provided in Appendix A.1.

**Figure 8:**
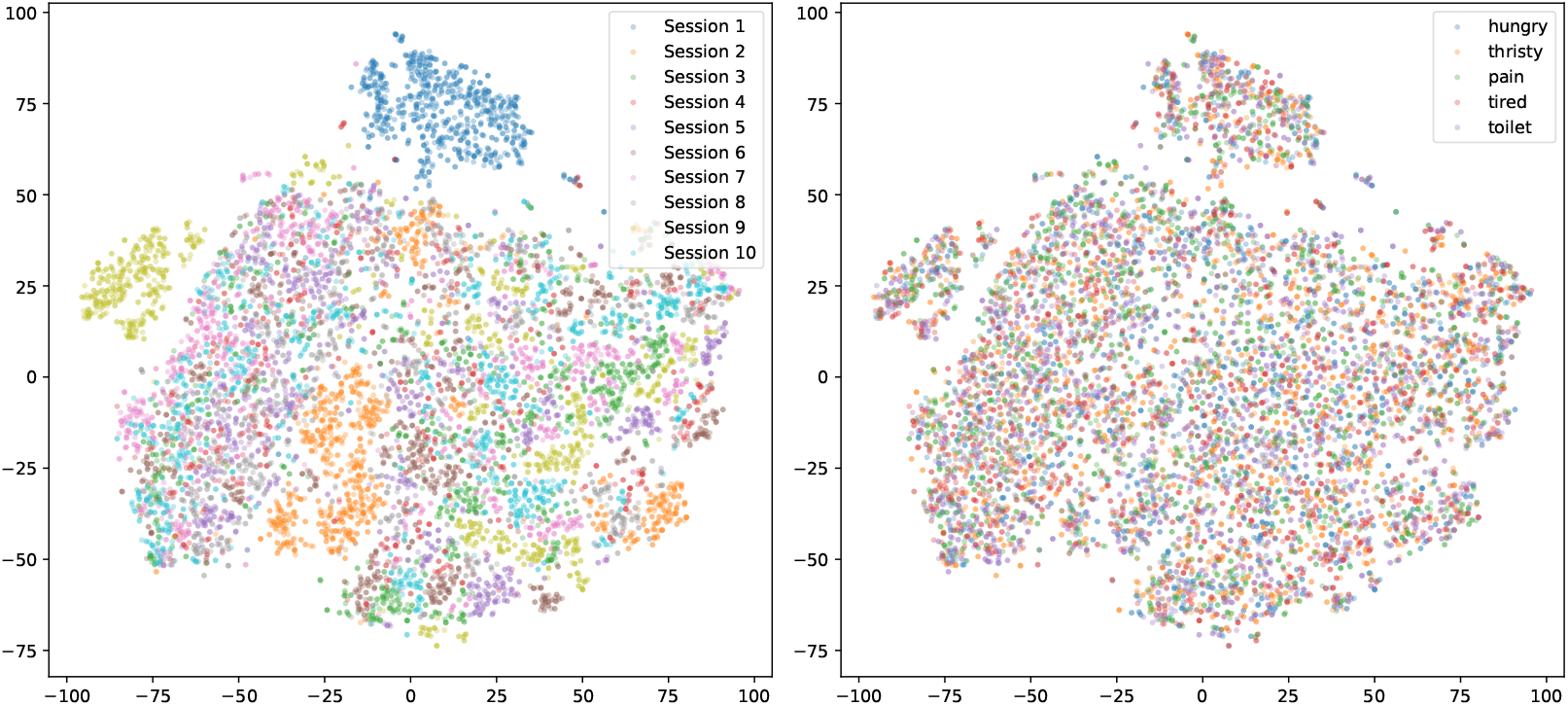
t-SNE projection of the per-trial covariances across the 10 EEG sessions of P4 in version 1. These are coloured according to the session label on the left, and according to the condition (word) on the right.

### 3.3 Decoding inner speech in experiment version 1

While we collected reading data in all experiments, we will only present results from version 2, since that version was specifically aimed at analysing the silent reading task. In this section we will first analyse the MEG data, followed by results from the 10 EEG sessions of P4.

We attempted methods such as sliding window LDA, full-epoch LDA, and the linear neural network. Running full-epoch models on the 1-second inner speech trials yielded chance-level results when decoding which word is being used in inner speech. We also attempted sliding-window decoding on a subset of MEG channels overlying the language area. However, the decoding accuracy timecourse exhibited substantial fluctuations and never exceeded 24%, where 20% is chance level. Thus, this is also a negative result.

On the EEG data of the generative inner speech trials of P4, LDA models were trained on each session utilising the channel-covariance over the 1-second epoch as features with 5-fold cross-validation. For this analysis preprocessing involved a 1-40 Hz bandpass filter and no ICA artefact removal was employed. Before computing covariance, trials were normalised to unit variance and zero mean. We found above 25% validation accuracy in only 3 sessions (Figure 9), with chance level being 20%. However, when correcting for multiple comparisons none of the sessions had significantly better performance than chance. It may be that by running more cross-validation folds the performance in some sessions reaches significance. Decoding the repetitive inner speech trials, or both types together did not produce better results.

**Figure 9:**
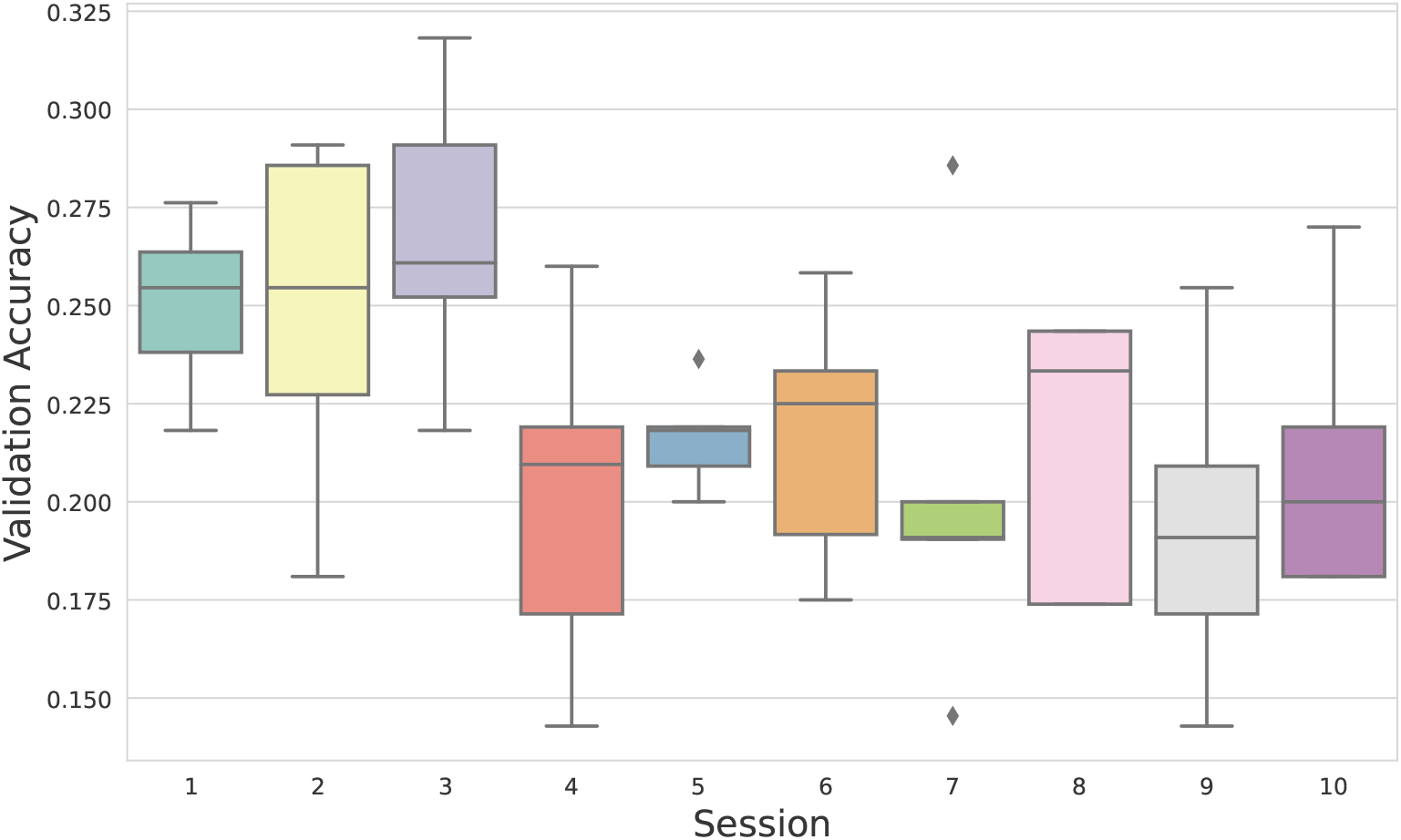
Validation accuracy distributions across the 5 folds of the 10 EEG sessions of P4 in experiment version 1. Separate LDA models are trained and evaluated on each fold and session to decode which of the 5 words is being used in the 1-second inner speech trials. Chance level is 0.2.

Nevertheless, we trained a single LDA model across the 3 best sessions, achieving 33% cross-validation accuracy. The same per-session folds were employed as in the previous analysis. To train a single LDA model across sessions, we made some modifications to the decoding pipeline. Rather than using the 1-second trial, we utilised the entire 4-second epoch with the four consecutive cues to compute covariance. To account for between-session differences, the mean session-level evoked response was subtracted from each trial before covariance computation, and the mean session-level covariance was also subtracted from each trial’s covariance. Since we evaluated many different methods on this data, and we selected these 3 sessions based on the previous analysis, there is a risk this result is inflated. Running the same decoding approach on all 10 sessions reduced cross-validated accuracy to 23.2%.

While the EEG data provides promising results, with some sessions exhibiting above-chance decoding accuracy, we must be cautious about drawing robust conclusions, due to the limited performance and risk of overfitting. The limited inner speech performance precluded assessment of transfer between silent reading and inner speech tasks. Evaluating transferability between sessions was also not feasible as most sessions displayed chance-level performance.

### 3.4 Decoding silent reading in experiment version 2

In this version, we collected a substantial number of silent reading trials only from 3 participants, across 4 modalities. For each session, we implemented an LDA model with 5-fold cross-validation (Figure 10). We utilised the 500 ms following word onset as our examples for decoding. For CTF, Elekta, and OPM data, the dimensionality of the LDA reduction was set to 50, and for EEG to 20.

**Figure 10:**
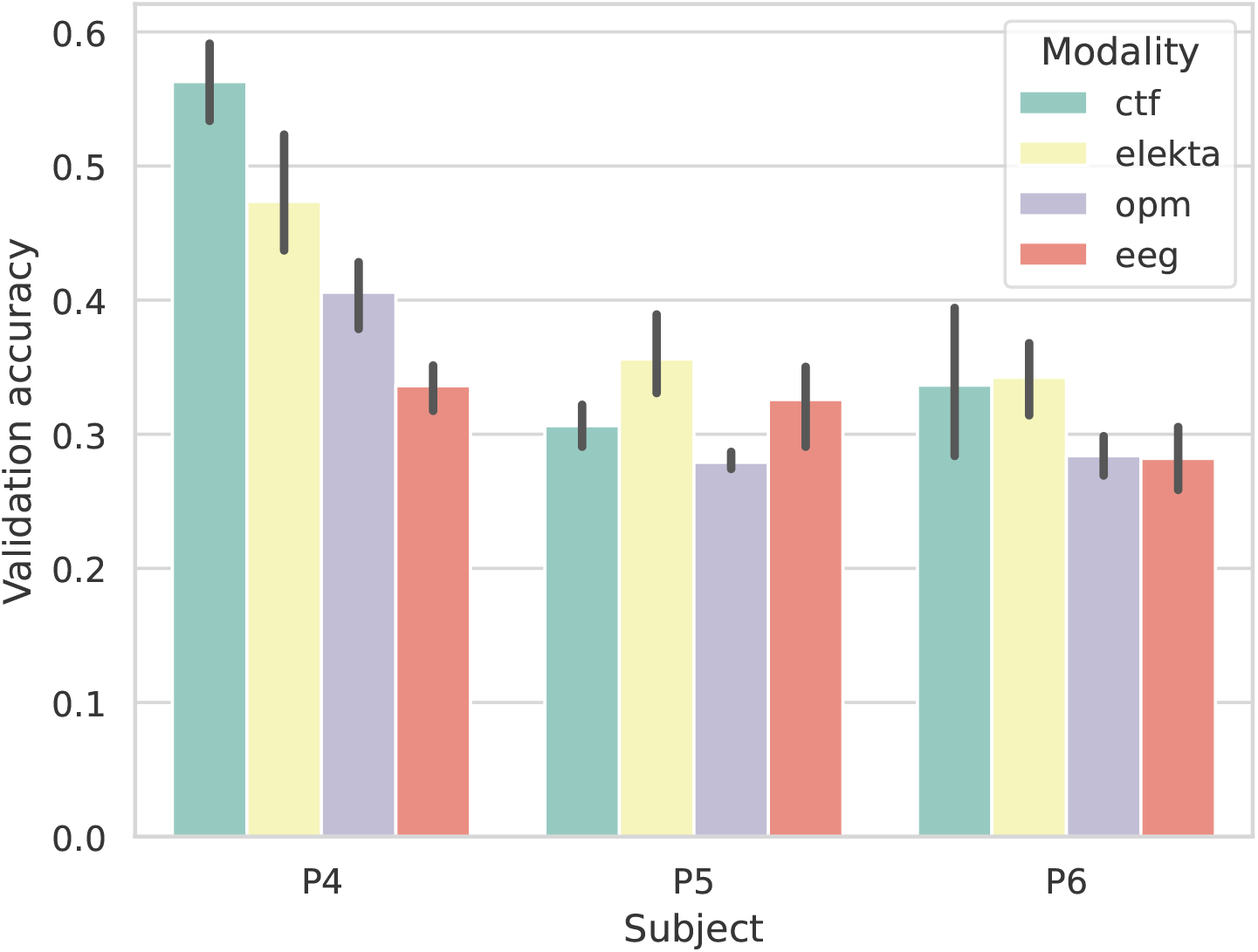
Validation accuracy (across 5 folds) for each session in experiment version 2. Separate LDA models are trained and evaluated on each fold and session to decode which word is presented during the 1-second trials. Black bars indicate 95% confidence interval. Chance level is 0.2 due to having 5 words with equal trial counts.

Accuracies are generally low, providing evidence that even with the visual component and numerous trials, silent reading is a challenging decoding task. P4 exhibits higher performance than the other two participants across all modalities except EEG. While CTF data achieved the best performance for P4, followed by Elekta, OPM and EEG, this is not the case for the other participants. Across modalities, P5 and P6 display more comparable performance. CTF and Elekta appear higher, while OPM and EEG are slightly lower but similar (especially for P6). The discrepancy between the CTF results for P4 and P6/P5 is particularly surprising. As depicted in Figure 4, unfortunately, due to experimental difficulties, the OPM sensor coverage of the visual area was inferior in these participants compared to P4. This could potentially explain the lower OPM performance.

It is difficult to derive conclusions from these results, and the experiment should be replicated across more subjects to enable a more robust comparison between modalities. It seems that traditional MEG scanners exhibit the best performance, while EEG and OPM lag behind but are comparable. This provides evidence that challenging decoding tasks such as silent reading are feasible with OPMs. Further innovation and better spatial coverage should enhance OPM decoding performance to approach traditional MEG.

Next, we investigated the temporal and spatial permutation feature importance (PFI) of the decoding models. We expect that PFI should appear similar across modalities. When plotting the spatial PFI for each modality we average across subjects and cross-validation folds. We utilised 20 permutations and set the number of nearby sensor locations for the spatial window to 4 in all modalities. Note this equates to 12 channels for Elekta, 8 channels for CTF, and 12 channels for OPMs, since these modalities contained multiple sensors at the same site. We depict the spatial PFI of Elekta, CTF, and EEG in Figure 11. It is clear that the visual area drives decoding across all modalities. We plot the OPM accuracy loss maps separately for each participant due to variability in available sensors (Figure 12). This demonstrates similar visual importance in P4 and P6. PFI was ineffective in P5, possibly due to limited decoding performance.

**Figure 11:**
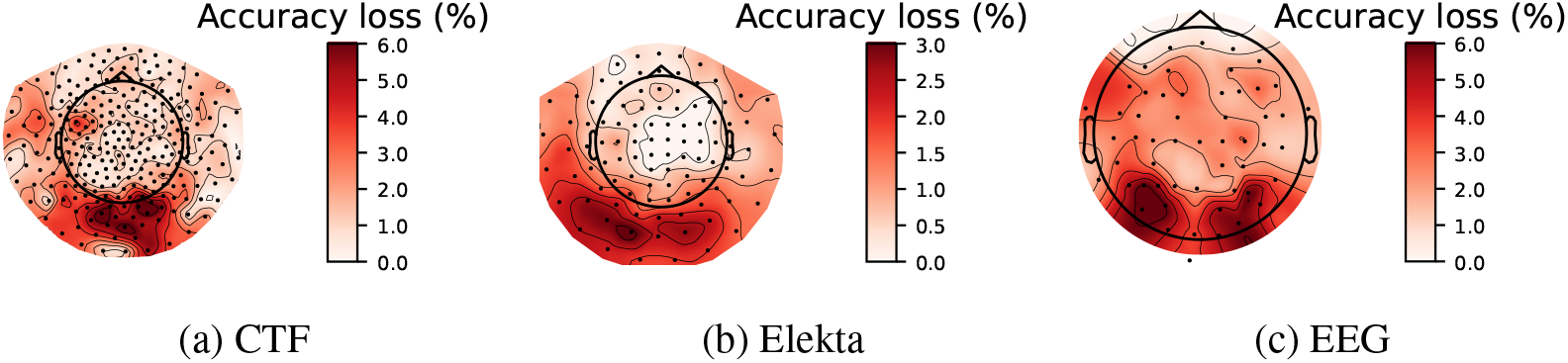
Sensor importance maps averaged across subjects for 3 modalities in experiment version 2. The importance maps are obtained by running spatial PFI on the trained LDA decoding models. Darker red shading indicates higher accuracy loss and thus higher stimulus-related information content.

**Figure 12:**
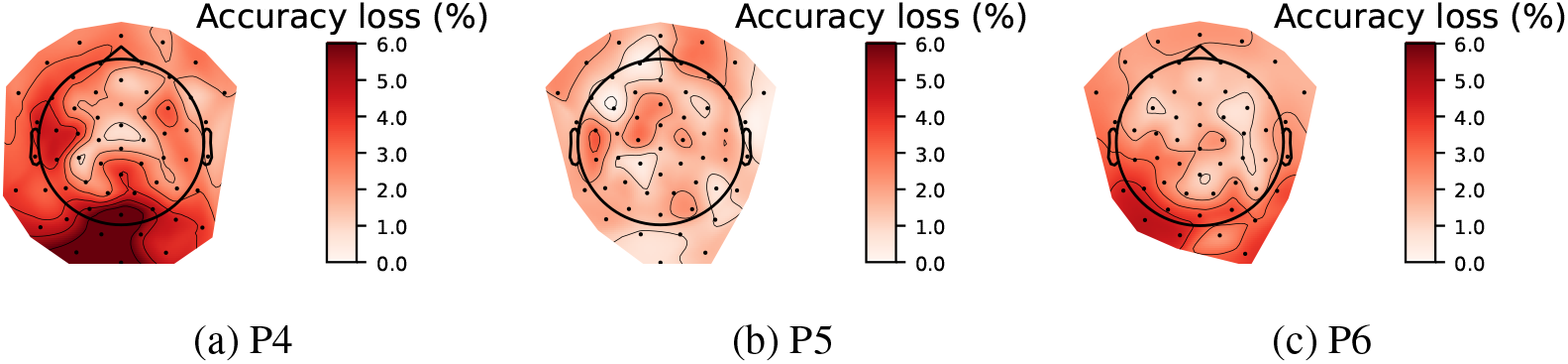
Sensor importance maps across subjects on the OPM recordings of experiment version 2. The importance maps are obtained by running spatial PFI on the trained LDA decoding models. Darker red shading indicates higher accuracy loss and thus higher stimulus-related information content. Note that P5 and P6 had less channels available, hence the smaller topographic map.

Finally, we illustrate temporal PFI across subjects and modalities in Figure 13. We utilised 20 permutations and a temporal window of 100 ms. Since decoding is driven by the visual region, we would anticipate peak accuracy loss around 150ms, which is indeed evident for P4. This subject also exhibits much less noisy timecourses. Across modalities, timecourses appear similar. Notably, all modalities except OPM display a second, smaller peak around 250ms. In other subjects, accuracy loss peaks later in the trial. This could reflect slower reaction times when silently reading. Interestingly, the CTF data for P6 and the EEG data for P5 exhibit two distinct peaks. This could indicate decoding is driven by both visual word processing and language processing due to silent reading. These plots highlight substantial between-subject variability in task-related brain-activity.

**Figure 13:**
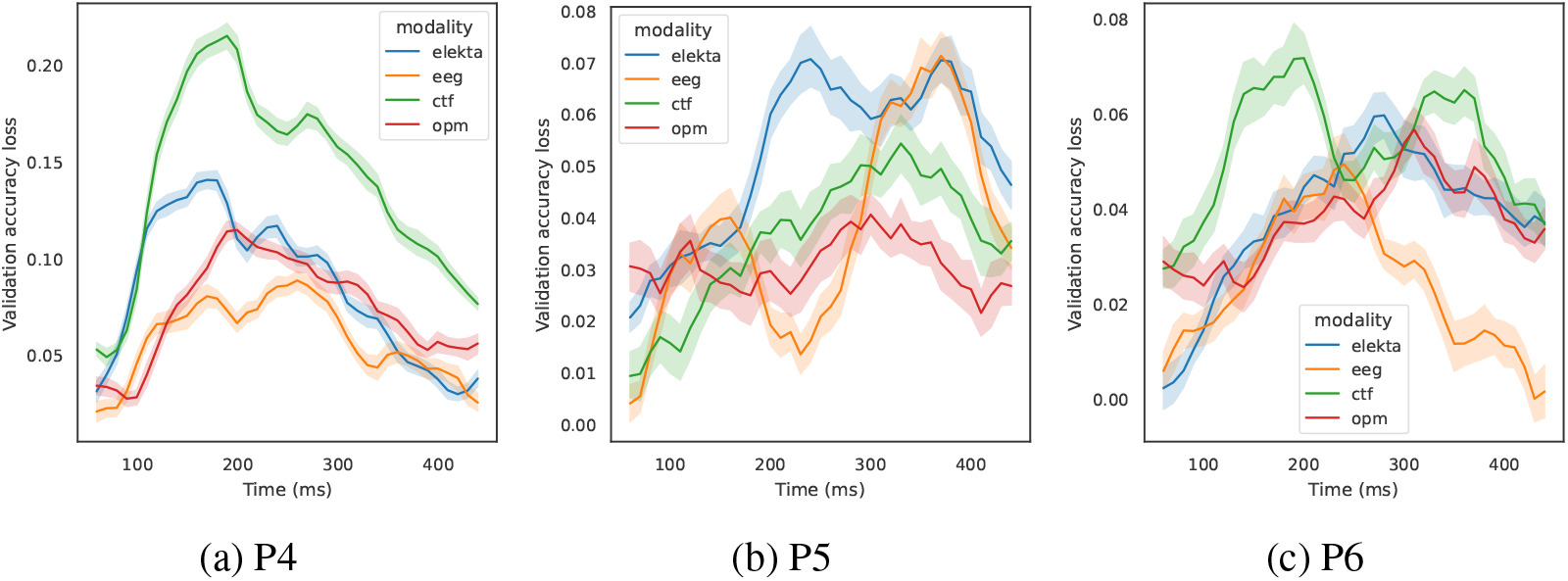
Temporal PFI across the 3 subjects (P4, P5, P6) and 4 modalities (lines with different colours) in experiment version 2. The timecourses are obtained by running temporal PFI on the trained LDA decodig models. Shading indicates 95% confidence interval across PFI permutations. The horizontal axis indicates time since stimulus onset.

## 4 Discussion

In this discussion we will first present and review related works for decoding inner speech from invasive and non-invasive modalities. Then in Section 4.5 we summarise our findings and conclude this chapter.

Typical experimental paradigms used in neuroimaging studies of inner speech include silent word repetition and silent reading, similar to ours. As reviewed by Geva (2018), these studies consistently demonstrate that compared to overt speech conditions, inner speech leads to reduced activation in motor regions like primary motor cortex and auditory sensory areas like primary auditory cortex. Inner speech also shows less engagement of sensory feedback regions like the superior temporal sulcus. However, inner speech robustly activates left hemisphere language regions. The authors suggest this enhanced activation in phonological and semantic regions may serve as an automatic compensatory mechanism that augments inner speech performance given the reduced sensory and motor feedback.

However, accurately interpreting the results of neuroimaging studies on inner speech remains challenging. Methodological limitations include controlling for overt speech production during inner speech tasks and ensuring participants are actually using inner speech rather than alternative cognitive strategies. Research on visual and motor imagery highlights that imagery across modalities relies on networks similar to actual perception and action (Kosslyn and Pylyshyn, 1994), reflecting shared neural processing. However, identifying a consistent substrate for imagery across all sensory modalities has proven more difficult.

When comparing invasive and non-invasive methods, the only major differences are in the brain signals and noise characteristics of each recording modality. The cognitive process of interest - inner speech itself - remains constant across invasive and non-invasive recordings. Invasive methods can record neural activity with much higher spatial resolution and signal-to-noise ratio (SNR). Thus, the key question becomes determining how reduced resolution and SNR in non-invasive EEG or MEG impacts detectability of inner speech processes. We discuss relevant invasive research of inner speech next.

### 4.1 Invasive methods

Inner speech has been most successfully studied through invasive methods. Electrocor-ticography (ECoG), where electrode arrays are placed below the skull directly on the brain surface, is one such technique. More invasive methods involve implanting electrodes within cortical tissue to record single neuron activity.

Martin et al. (2016) demonstrate one of the first successful decodings of individual words during imagined speech from direct cortical recordings in humans. Their study design involved recording high gamma activity using ECoG during listening, overt speech, and imagined speech conditions for 6 different words. They developed a binary classification approach using support vector machines that incorporated dynamic time warping (DTW) to account for temporal variability in speech production. At the group level, classification accuracy was significantly above chance for imagined speech, with the best word pair reaching 88% accuracy. However, across all word pairs accuracy was much lower, and performance was variable between subjects. Discriminative information was located primarily in the superior temporal gyrus, inferior frontal gyrus, and sensorimotor cortex, consistent with their role in speech processing (Hickok and Poeppel, 2007). This work can inform future noninvasive research on which brain areas to focus on and employ techniques like DTW to handle temporal variability.

Recent work by Wandelt et al. (2022) demonstrates the feasibility of decoding internal speech from single neuron activity in the supramarginal gyrus (SMG) and somatosensory cortex (S1) of a tetraplegic human participant. Their study design allowed comparison of neural activity between visual word reading, listening, vocalised, and inner speech across 8 words. The authors found individual SMG neurons showed selective tuning to specific words during the internal speech condition. Using these neural signals, they achieved up to 91% decoding accuracy for internal speech words using a real-time setting.

Importantly, the authors found strong shared neural representations in SMG between internal speech, reading visually presented words, and vocalised speech production. This points to the involvement of common underlying cognitive processes between these tasks. Their decoder was also robust to different internal speech strategies, such as auditory vs. visual imagery, suggesting flexibility for individual mental strategies in future applications. Specifically, they tested the generalisation performance of the decoder between all tasks. Decoders trained on auditory cue trials were less generalisable to inner and vocalised speech than those trained on written cue trials. This shows silent reading brain activity may be closer to pure inner speech. While the cue modalities were separable during the cue-phase brain activity, they overlapped during subsequent phases. Thus, internal and vocalised speech representations may not be influenced by the cue modality, a promising result for repetitive inner speech paradigms.

In a different line of work, Willett et al. (2021) demonstrate the potential for real-time decoding of attempted handwriting movements from neural activity as a high-speed BCI. In this study, a participant with tetraplegia from spinal cord injury attempted to handwrite letters and words by imagining holding a pen and writing. Neural activity was recorded from intracortical electrode arrays implanted in the hand area of motor cortex. They found individual neurons showed selective patterns of activation for different handwritten letters, enabling reconstruction of pen trajectories. While not direct inner speech, it remains a purely imagined task with no external stimuli or produced behaviour. In online experiments, the participant achieved remarkable typing speeds of over 90 characters per minute, with over 94% raw accuracy. While an invasive approach was used, the neural dynamics revealed in motor cortex potentially inform non-invasive BCI design. Attempted handwriting may be a promising paradigm for EEG and imaging BCIs if cortical patterns can be sufficiently resolved.

While limited in humans for ethical reasons, invasive recordings in neurological patients provide unparalleled detailed characterisation of the neurophysiology underlying inner speech phenomena. These studies unequivocally show that inner speech and movement imagery decoding are possible invasively, and demonstrate their potential in BCI applications. Next, we turn to non-invasive studies of inner speech.

### 4.2 EEG

While invasive studies are rare due to the nature of intracranial recordings, EEG studies of inner speech are also uncommon because of the difficulty in overcoming the low signal-to-noise ratio and spatial resolution inherent in scalp EEG recordings.

Cooney et al. (2019) investigate using CNNs to classify imagined spoken word-pairs from EEG signals. Their dataset contained 6 Spanish words imagined by 15 subjects. All 15 possible word-pairs were extracted and EEG signals corresponding to an early imagined speech time-window were used. Results showed a deep CNN achieved the best average accuracy of 62.37% across subjects and word-pairs. This performance however is still barely above chance level, indicating the ongoing difficulty of decoding imagined speech from noisy EEG recordings.

Ling et al. (2019) investigate how visual words are represented in the brain using EEG-based decoding and image reconstruction techniques. Their study had 14 participants view 80 high-frequency nouns while recording EEG data. They then used multivariate pattern analysis on the EEG data to decode visual and orthographic properties of the words. Specifically, they were able to decode pairwise word discriminability well above chance across participants, with peak performance around 170ms after stimulus onset. They also applied representational similarity analysis to show the word decoding results correlated with visual and orthographic similarity but not semantic similarity. This is perhaps unsurprising as decoding visual activity is well-studied with EEG.

### 4.3 MEG

Défossez et al. (2022) present a novel method for decoding natural continuous speech from non-invasive MEG and EEG recordings. The authors leverage recent advances in self-supervised speech representation learning, specifically wav2vec 2.0 (Baevski et al., 2020), to obtain semantically meaningful speech embeddings from raw audio. These speech embeddings are aligned with M/EEG signals recorded while participants passively listened to audio samples. A joint CNN architecture with a contrastive loss is used to predict the speech embeddings from the neural signals. Without any individual calibration, their model can identify 3-second speech segments with up to 72.5% top-10 accuracy across nearly 1,600 samples for MEG and 19.1% across 2,600 samples for EEG.

For decoding inner speech, this study provides a promising framework to handle individual variability and extract meaningful speech features from limited data. The zero-shot decoding is particularly impressive, as it avoids constraints of classifiers trained on small stimulus sets. However, additional work is needed to apply this to inner speech due to the lack of audible signals for alignment. They leverage both group modelling and training large models across multiple datasets. Their approach is well suited to incorporating forecasting or other self-supervised objectives. The contrastive loss allows for out-of-distribution decoding, as it is not limited by the categorical nature of standard classifiers. Furthermore by incorporating multiple feature extractors (e.g. CNNs trained on images), the same contrastive approach and brain model can be applied to various decoding tasks.

Direct MEG investigations of inner speech are limited. Dash et al. (2020a) decode 5 imagined and overtly spoken phrases from MEG. Three decoding methods were tested: an artificial neural network (ANN) using statistical features, a CNN on time-frequency images, and a CNN with combined spatial, spectral and temporal features. The CNN approaches significantly outperformed the ANN, achieving up to 96% accuracy for spoken phrases and 93% for imagined phrases with the combined features. A key limitation is using phrases which likely contain more decodable information but are harder to scale up. A contrastive approach, or decoding at the phoneme/word level is more desirable.

### 4.4 OPM-MEG

While superconducting quantum interference devices (SQUIDs) are traditionally used for MEG, optically pumped magnetometers (OPMs) have recently emerged as a promising alternative for MEG measurements (Boto et al., 2018). OPMs offer several advantages over SQUIDs including room-temperature operation, lower cost, higher sensitivity, and allow head motion (Boto et al., 2017). Their compact size also enables flexible sensor arrays that can be customized to target specific brain regions or conform to individual head shapes (Boto et al., 2018).

A key application of MEG is non-invasive decoding of mental states and cognitive processes from neural activity patterns. Some initial studies have now explored using OPM-MEG for neural decoding. Wittevrongel et al. (2021) demonstrate OPM-MEG enables robust singletrial analysis and real-time decoding for BCI applications. They compared OPM-MEG and EEG for decoding visual evoked responses, including event-related potentials/fields (ERPs/ERFs) to motion-onset and steady-state visual evoked potentials (SSVEPs) to flickering stimuli. For motion-onset, OPM-MEG and EEG showed similar ERP/ERF components (N/M200, P/M300) with comparable signal-to-noise ratios. For SSVEPs, OPMs had higher SNR in the high frequency range (25-29 Hz) while EEG was better at low frequencies (8-12 Hz). In a real-time SSVEP spelling task, OPM-MEG achieved 97.7% average accuracy comparable to state-of-the-art EEG systems. These results validate OPM-MEG for robust single-trial decoding in BCI applications. The improved SNR and spatial resolution suggest OPMs could enable more advanced decoding capabilities. Their wearable and flexible nature makes them well-suited for practical BCI applications.

### 4.5 Conclusion

This chapter presented an in-depth investigation into decoding inner speech and silent reading from non-invasive electrophysiological recordings. The key findings were the following. Silent reading of words could be decoded from EEG, MEG, and OPMs with 30-40% accuracy across 5 words, driven by early visual processing. Comparing modalities showed traditional MEG had the best performance for decoding silent reading, while OPMs and EEG had lower but comparable accuracies. Inner speech decoding was mostly at chance levels in EEG and MEG across multiple decoding approaches. The highest accuracy reached for inner speech was 33% across 3 EEG sessions using covariance features, but the validity of this result is debatable.

Our silent reading results demonstrate the feasibility of decoding visual representations of words from non-invasive recordings, consistent with prior EEG and MEG decoding studies (Chan et al., 2011; Ling et al., 2019). The decoding appeared to be driven by early visual responses, with a later peak potentially reflecting higher-level language processing (Kutas and Van Petten, 1988). This late component merits further investigation as a marker of semantic processing. While more subjects are needed for a robust comparison, OPMs achieved decoding accuracy comparable to that of EEG. In one participant with good spatial coverage, OPM decoding performance was even better than EEG, highlighting their promise given advantages like wearability. Dense coverage of visual regions may be critical for the investigated decoding task. Our results also underscored the high between-subject variability, both in overall performance and in the timing of informative decoding features.

In contrast to silent reading, our extensive efforts to decode two types of inner speech were largely unsuccessful across EEG and MEG. While we explored various decoding algorithms and experimental designs, accuracy never substantially exceeded chance levels. This contrasts with more promising results from intracranial recordings in humans (Martin et al., 2016; Wandelt et al., 2022), and suggests non-invasive signals may not adequately capture the subtle dynamics of inner speech. There was also substantial between-session variability.

In addition to the analyses presented, numerous unsuccessful decoding approaches were pursued on the inner speech data. On the MEG recordings, these included logistic regression, CNNs, SVMs, concatenating or averaging consecutive trials, and per-session versus aggregated-session decoding. For EEG, besides the MEG-based methods, other unsuccessful attempts involved temporal alignment of trials, PCA denoising, Riemannian classifiers, baseline correction, and wavelet features. We tried several referencing approaches such as common average reference, mastoid references, and current source density estimation through the Laplacian method. We also tried wider bandwidth filters (up to 100Hz), however we noticed no improvement with the addition of gamma activity, even though it is reported in invasive inner speech studies. Noise and superposition in scalp measurements may overshadow weak gamma oscillations.

Several factors could underlie the difficulty of decoding inner speech non-invasively. Inner speech lacks the external stimuli and muscle activations present during overt tasks, reducing the signal-to-noise ratio. There is also high inter-individual variability in inner speech strategies. Here we focused on collecting large trial counts from a few participants rather than a small sample across many subjects. Our cross-cue paradigm may also induce visual confounds that overshadow inner speech signals. Having participants repeatedly imagine brief, single words likely differs from natural inner speech involving longer phrases. Further limitations of our work include the small number of participants and the small set of words.

Future investigations could explore alternative paradigms more representative of natural speech, such as imagining longer phrases or reading whole sentences silently. Transfer learning and self-supervision may help extract robust inner speech representations amidst noise (Défossez et al., 2022). Intracranial findings point to superior temporal, inferior frontal, and motor areas as promising decoding targets. For non-invasive BCIs, approaches beyond word-level decoding may be needed for inner speech-based communication, such as decoding phonemes, or imagined handwriting.

One downside of our task design is that the low-level characteristics of the visual appearance of the words might be a confounding factor in our silent reading results. Future work should consider using varied graphical representations of the same word across trials.

In summary, our results highlight the significant challenges in decoding inner speech correlates non-invasively compared to overt tasks. Substantial innovation in experiments and analyses will likely be essential to enhance the fidelity of decoded inner speech for BCIs. While current decoding performance was limited, our proof of concept work provides a useful platform with extensive trial counts for future efforts at modelling inner speech. Having multiple sessions allows for testing across-session generalisability. Neuroscientific understanding of inner speech may be deepened through comparing the different experimental paradigms.

## A Supplementary Material

### A.1 Evoked analysis

We computed evoked responses jointly across the two inner speech types without any baseline correction. We visualise these for each session for electrode PO7 (visual area) in Figure 14. It is evident that evoked responses across sessions are very similar in the visual area, except for session 2 which appears to be an outlier. The plot also shows the expected response to visual stimulus, with the first peak as early as 100 ms post-stimulus (P100), followed by several peaks and troughs. This demonstrates the oscillatory nature of the evoked response in the visual area, likely due to the cross cue used in the inner speech task.

**Figure 14:**
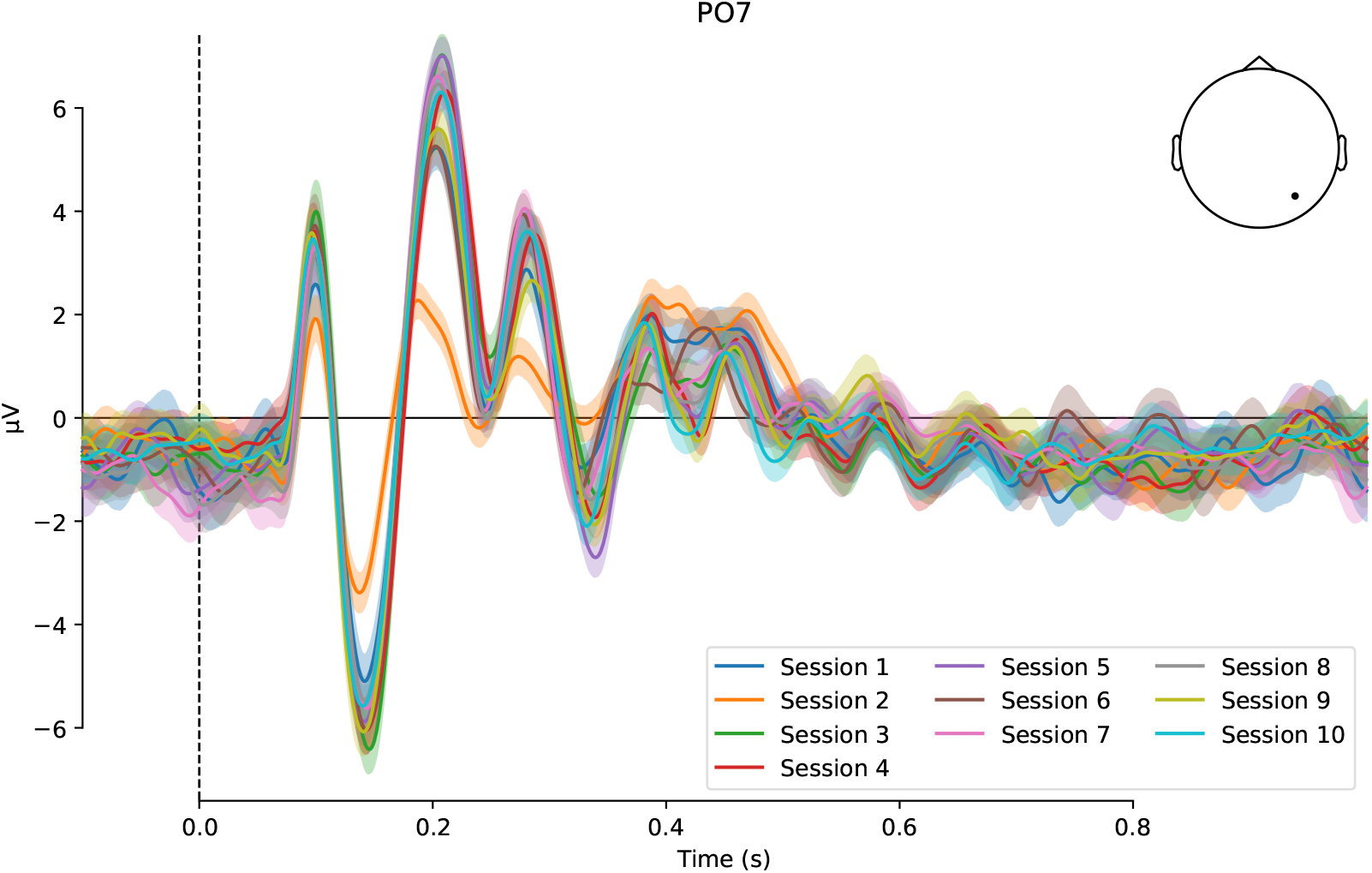
Evoked responses across the 10 EEG sessions of P4 for 1 electrode (PO7) in the visual area. Shading indicates 95% confidence interval across trials. Timepoint 0 indicates stimulus (cross) onset.

While we were also interested to compare other channels across sessions, non-visual channels exhibit more noise. Thus, we plot evoked responses in separate plots per-session. Figure 15 shows this for the T7 electrode, which is above the temporal lobe. Evoked responses in the temporal lobe are much more variable across sessions, but all display peak activity around 400 ms post-stimulus. However, it is questionable whether this reflects language-related activity (due to inner speech), or merely spreading/propagation of the visual response. The latter is more probable.

**Figure 15:**
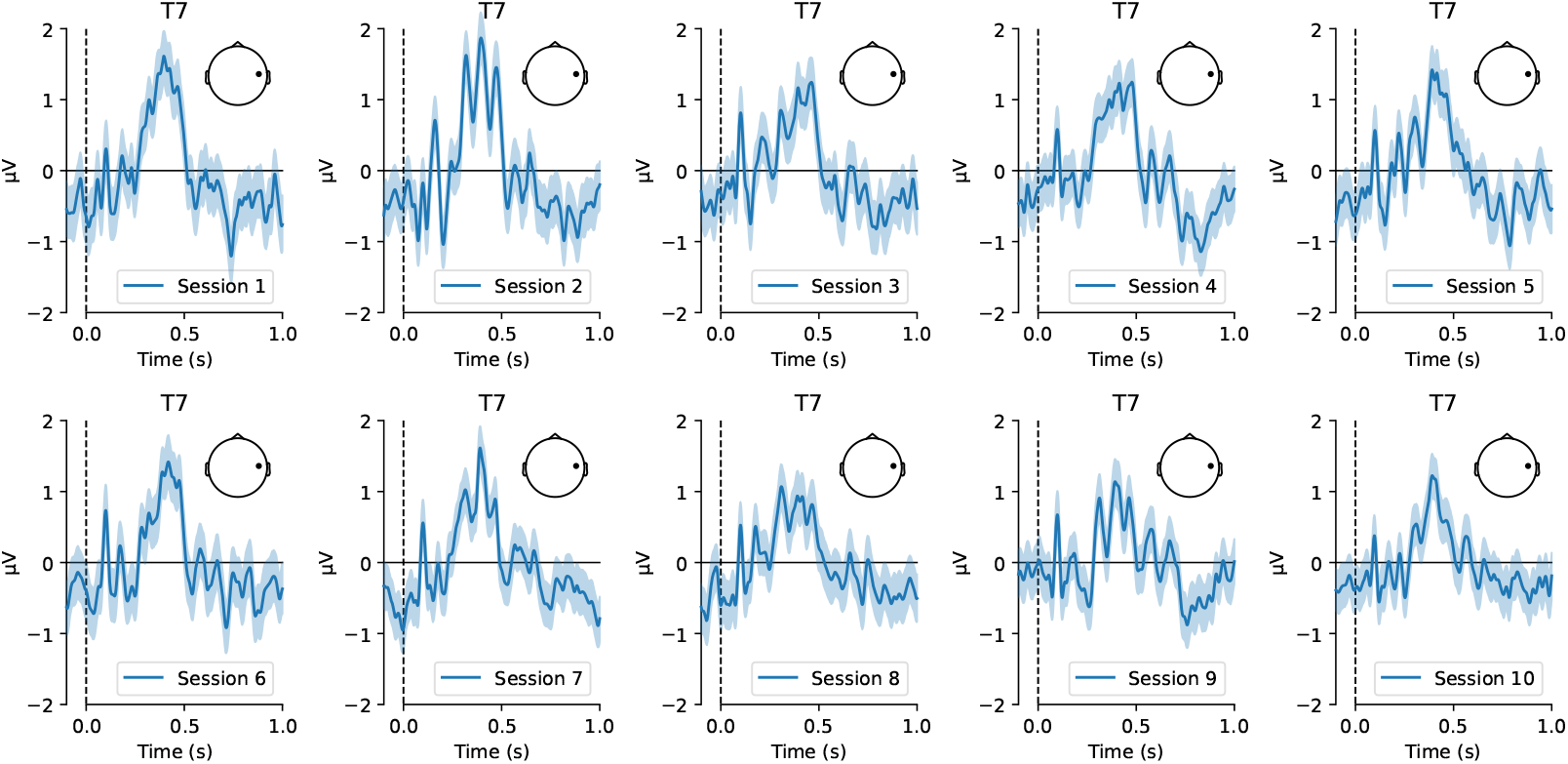
Evoked responses across the 10 EEG sessions of P4 for 1 electrode (T7) above the temporal lobe. Shading indicates 95% confidence interval across trials. Timepoint 0 indicates stimulus (cross) onset.

It is important to note that since we utilise the Cz electrode for referencing, our evoked results are influenced by this. Any evoked response present at the reference is subtracted from all other channels. To better elucidate the spatiotemporal evolution of the evoked response, we plot responses averaged across sessions for all channels concurrently (Figure 16). This demonstrates that after the initial two visual peaks, at 200 ms a third positive visual peak emerges, accompanied by a smaller negative activity in the frontal area. Then, at 285 ms another smaller visual peak occurs, followed by a negative peak at 334 ms. At this time, some positive activity also arises in the frontal area. Finally, more visual positivity is observed around 386 ms, which shifts slightly to temporal/lateral areas at 406 ms, returning to the visual area at 452 ms. This plot provides a robust characterisation of the evoked response to inner speech across a substantial number of trials and sessions. However, we suspect the described activity remains principally due to the cross-cue presentation.

**Figure 16:**
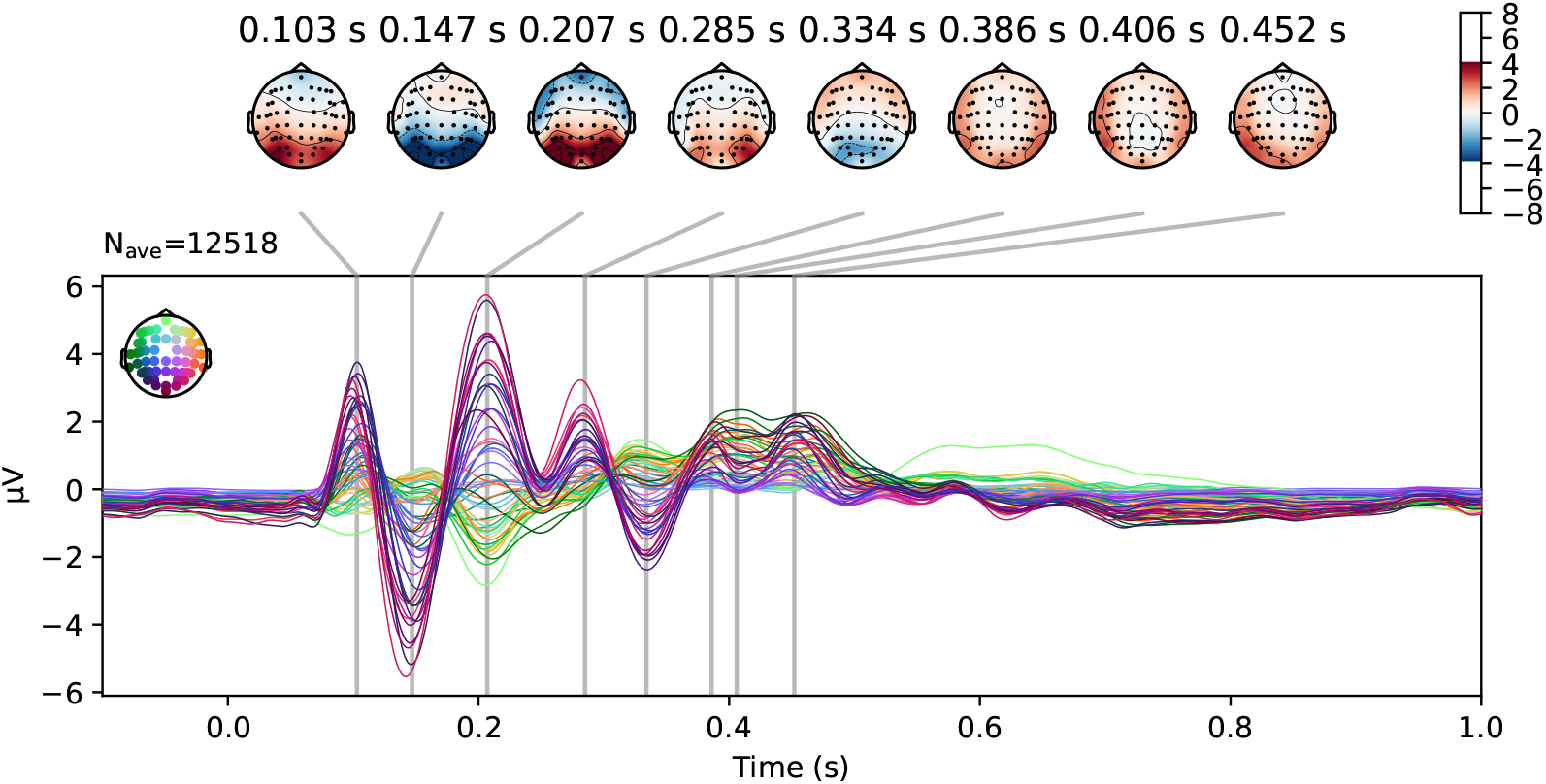
Joint evoked responses for each channel averaged across all 10 EEG sessions of P4. The spatial topography and timestamp of notable peaks is shown in the upper part.

Figure 17 displays the evoked response for each word across all sessions. Again, we only examine the two inner speech types here. We also opted to plot the entire 4-second trial with the 4 consecutive crosses, rather than averaging over these. We can discern that as the trial continues, the response to subsequent cross cues diminishes. This could reflect genuine activity changes and/or more noise across sessions later in the 4-second trial. There are no apparent differences between the evoked responses of words. This is anticipated since inner speech should elicit very subtle distinctions in EEG that would be nullified when averaging over many trials and sessions.

**Figure 17:**
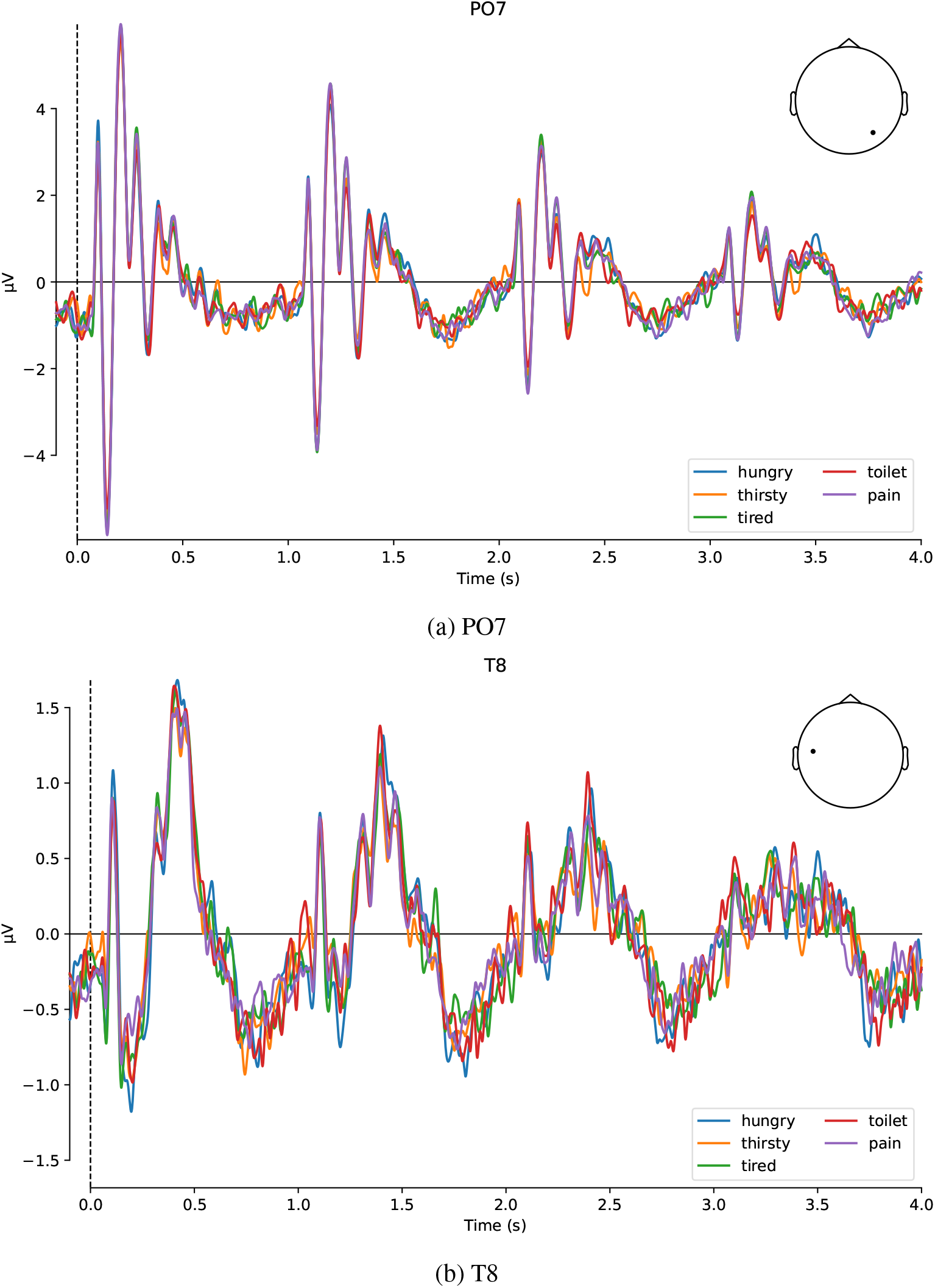
4-second evoked responses in 2 channels (PO7 and T8) across the 5 words averaged across all 10 EEG sessions of P4. Each line represents a different word.

We wanted to verify that the evoked responses are purely visual in nature. However, there is no straightforward way to separate the visual and inner speech-related activity. The cross cue is essential to provide consistent timing for inner speech production; otherwise, variability in timing would impede decoding. Still, we conducted 1 EEG session with P4, where we implemented 3 tasks. First, we utilised the standard repetitive inner speech task with the 4 consecutive cues (*cue+inner speech*). Then, we included a task where the visual stimuli were identical, but the participant was instructed not to think/internally vocalise the words, simply observe the crosses (*cue-only* task). Finally, we incorporated a task with only 1 cross cue at the beginning of the 4-second trial, after which the participant attempted to repeat the inner speech 4 times as in the original task, but without timing alignment (*inner speech-only* task).

Evoked responses across the 4-second trials for the three tasks from this single session are depicted in Figure 18. This provides unambiguous evidence that the previously observed evoked responses are elicited by the cross cue. There are no discernible differences between the brain activity of solely observing the cues, compared to also engaging inner speech. The task where inner speech had to be repeated without visual cues shows that after the evoked response to the initial cue, there are no subsequent evoked responses attributable to inner speech. This could also stem from variability in timing. However, it is more likely that utilising inner speech is too subtle to generate brain signals exceeding baseline noise. These findings imply that decoding inner speech may be an equally challenging endeavour.

**Figure 18:**
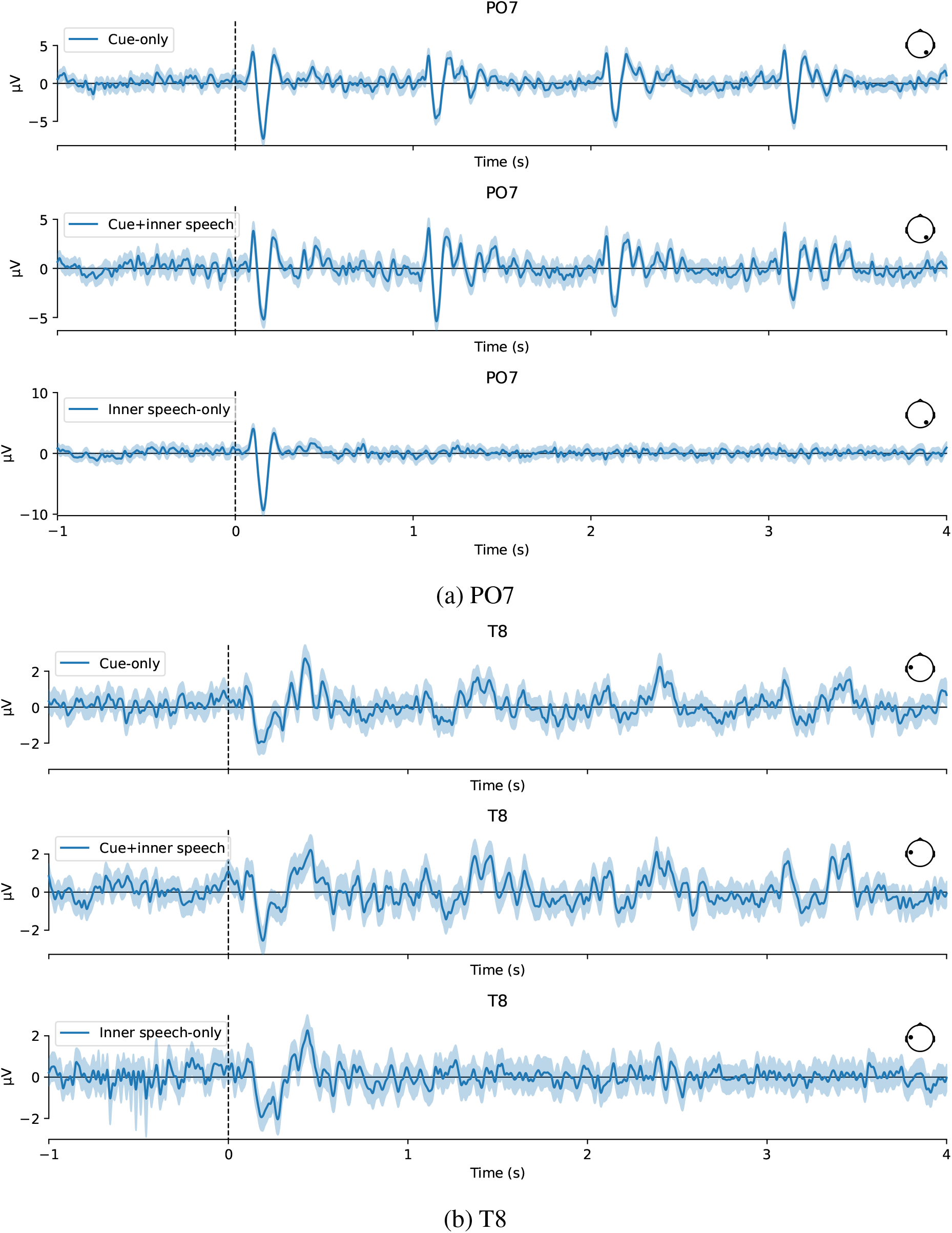
4-second evoked responses in 2 channels, PO7 and T8, for the EEG session with 3 tasks. The evoked response across the 4-second trial is shown for the cue-only (top), cue+inner speech (middle), and inner speech-only (bottom) tasks, in both (a) and (b). Shading indicates 95% confidence interval across trials.

## Footnotes

1 https://www.nottingham.ac.uk/research/groups/spmic/facilities/ctf-meg-scanner.aspx

2 https://github.com/OHBA-analysis/oslpy

## Notes

### Competing Interest Statement

The authors have declared no competing interest.

